# A self-supervised DNA foundation model with collapse-resistant multimodal fusion

**DOI:** 10.64898/2026.08.19.745697

**Authors:** Yuyan Chen

## Abstract

Genomic foundation models pretrained on DNA sequence have achieved strong performance across many tasks, but sequence-only representations cannot fully capture regulatory information from additional DNA-centric modalities. Existing multimodal genomic models are optimized for specific prediction tasks rather than reusable embeddings. Directly fusing heterogeneous modalities is challenging because sparse, peak-shaped regulatory signals and dense sequence embeddings have markedly different statistical structures, making naive alignment prone to near-zero solutions. We present a self-supervised DNA-centric multimodal foundation model integrating DNA sequence embeddings with local and global chromatin accessibility in a shared encoder to produce reusable window-level embeddings. We show that global normalization alleviates this collapse, enabling effective joint learning. The resulting embeddings improve regulatory activity prediction, regulatory signal ranking and chromatin accessibility peak detection, achieving a 4.6-fold AUPRC improvement over the DNA-only baseline, with further gains on external ClinVar, GTEx eQTL and PBMC caQTL datasets.

## Introduction

Modern genomic models increasingly aim to learn general representations of genomic regions from DNA sequence. DNA sequence contains fundamental regulatory information, including transcription factor motifs, conserved elements and long-range genomic context. However, many functional genomics tasks require information beyond sequence alone: regulatory activity prediction can benefit from chromatin accessibility and histone-mark signals^17,18^, variant-effect assessment often requires linking sequence variation to regulatory state and functional consequence, and expression-related analyses may depend on the relationship between regulatory elements and gene activity. Additional DNA-centric modalities, including chromatin accessibility, epigenomic marks, DNA methylation and expression-associated signals, provide complementary views of genome function. Integrating multiple DNA-centric modalities into genomic models can therefore produce representations that better capture regulatory function and support downstream analyses across chromatin-associated activity, gene regulation and variant effects.

Recent genomic models have developed along several complementary directions. DNA language models, including DNABERT-2, Nucleotide Transformer and HyenaDNA^1,2,3,19^, learn sequence-level representations from nucleotide or k-mer inputs, while long-range sequence models such as Enformer and AlphaGenome^4,5,20^ extend sequence modelling to distal regulatory context and functional prediction. Large-scale sequence models such as Evo 2^6^ further expand the capacity of sequence-only modelling. In parallel, chromatin accessibility and epigenomic models, including ChromBPNet, gReLU and EpiBERT^7,8,9^, have shown that regulatory signals provide rich information for modelling genome function. These advances have established the value of both sequence-derived and regulatory-state information, but most existing models still use these signals to optimize predefined prediction targets or modality-specific objectives. What remains underdeveloped is a DNA-centric multimodal embedding framework that integrates heterogeneous genomic signals into a shared representation, learns their complementary information and cross-modal interactions, and remains reusable across downstream tasks.

A key challenge is how to fuse heterogeneous genomic modalities into a stable shared representation. DNA-derived embeddings are dense and continuous, whereas regulatory signals such as chromatin accessibility are sparse and peak-shaped, with strong signals concentrated in a small fraction of positions. This distributional mismatch makes naive multimodal fusion prone to modality imbalance: the model may learn modality-specific shortcuts rather than meaningful cross-modal interactions. In self-supervised training, this imbalance can manifest as near-zero collapse of the sparse regulatory modality, limiting effective alignment between sequence-derived and regulatory information. Overcoming this failure mode is therefore critical for learning reusable DNA-centric multimodal embeddings from heterogeneous genomic signals.

Here, we develop DNA-MFM, a self-supervised DNA-centric multimodal foundation model for learning reusable region-level embeddings from heterogeneous genomic signals. The framework integrates DNA sequence embeddings with local and global chromatin accessibility signals into a shared representation space, allowing sequence-derived and regulatory information to be jointly encoded within the same genomic window. To make this fusion effective, we diagnose a sparse-modality collapse problem that arises when peak-shaped regulatory signals are directly aligned with dense sequence representations, and introduce a global-normalization strategy that stabilizes joint multimodal learning. Compared with DNA-only and existing sequence-based models, the resulting embeddings achieve stronger performance across three evaluation categories: pre-training reconstruction quality, assessed by three-modality masked reconstruction; internal regulatory-function benchmarks, including regulatory activity classification, regulatory activity ranking, chromatin peak detection and peak-position classification; and external functional and variant-effect validation, including ClinVar non-coding variant pathogenicity assessment, GTEx eQTL effect-size prediction and PBMC caQTL effect-size prediction. Systematic ablations further show that multimodal fusion, global normalization and embedding readout design each contribute to the final representation.

## Results

### A DNA-centric multimodal workflow for reusable genomic embeddings

**Fig. 1.**
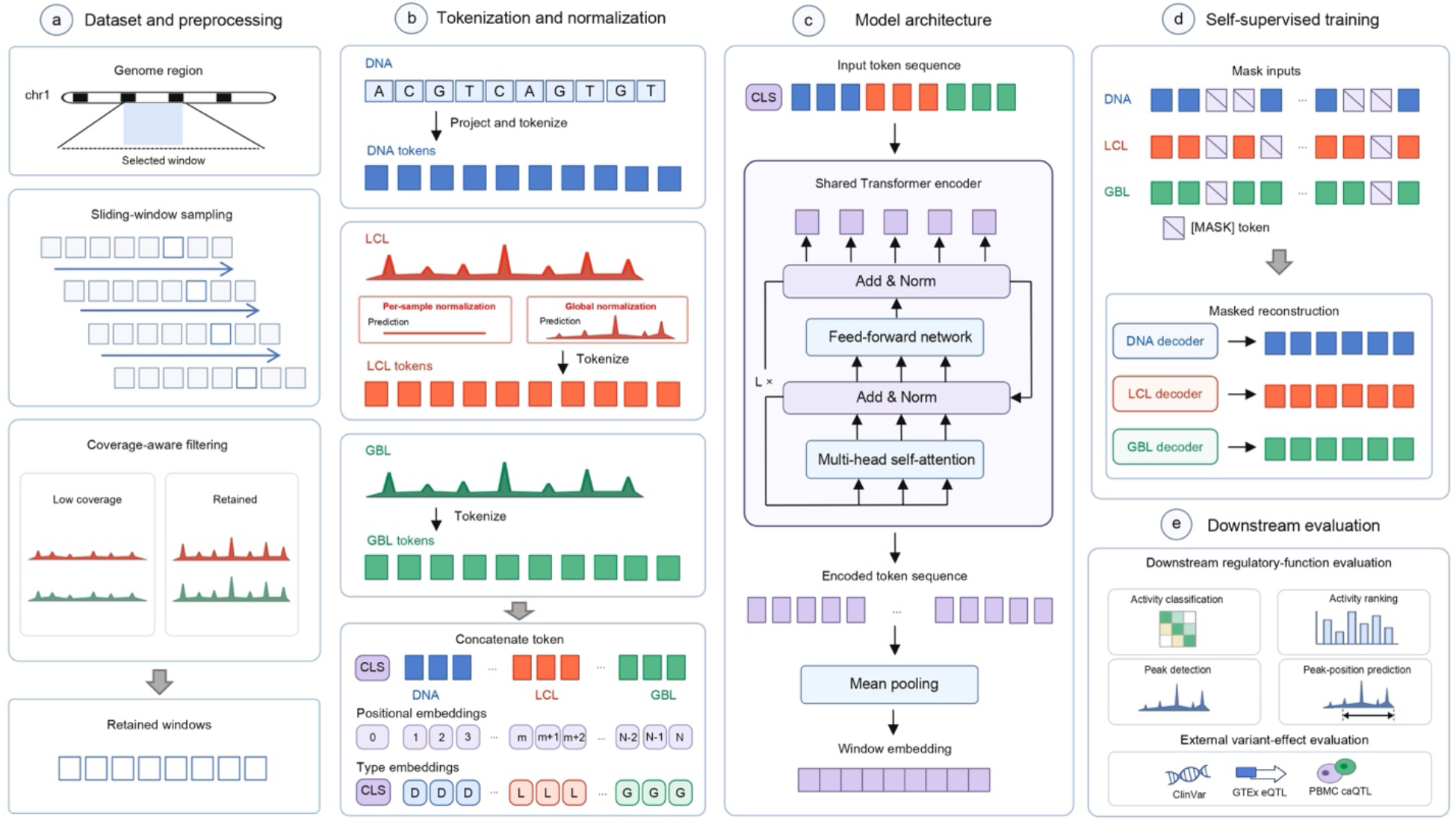
End-to-end workflow of the DNA-centric multimodal foundation model. (a) Genomic windows are constructed from the human reference genome with matched regulatory signals, using chromosomes 1-19 for pre-training and chromosomes 20-22 for held-out evaluation. (b) Each window is represented by DNA sequence embeddings, local chromatin accessibility and global chromatin accessibility. (c) The three modalities are normalized, tokenized and aligned in a shared representation space, with global normalization used to stabilize sparse accessibility signals. (d) The aligned representation is trained with modality-specific masking and continuous reconstruction. (e) The trained model generates reusable window-level embeddings for internal regulatory-function tasks and external ClinVar, GTEx eQTL and PBMC caQTL validation.

We first established DNA-MFM, an end-to-end DNA-centric multimodal foundation model for learning reusable genomic-window embeddings (Fig. 1). The workflow starts from genomic windows paired with matched regulatory signals, with each window represented by three co-registered streams from the same genomic region: DNA sequence embeddings, local chromatin accessibility and global chromatin accessibility. These heterogeneous inputs are normalized, tokenized and aligned within a shared representation space, allowing sequence-derived and regulatory information from the same window to be jointly encoded. To stabilize this fusion, the workflow incorporates global normalization for sparse, peak-shaped accessibility signals, preventing near-zero collapse during multimodal alignment. The model then produces a single window-level embedding that can be used for pre-training reconstruction, internal regulatory-function benchmarks and external functional or variant-effect validation. This overview defines the common representation pipeline used throughout the following experiments.

### Modality-specific behavior of masked reconstruction

**Fig. 2.**
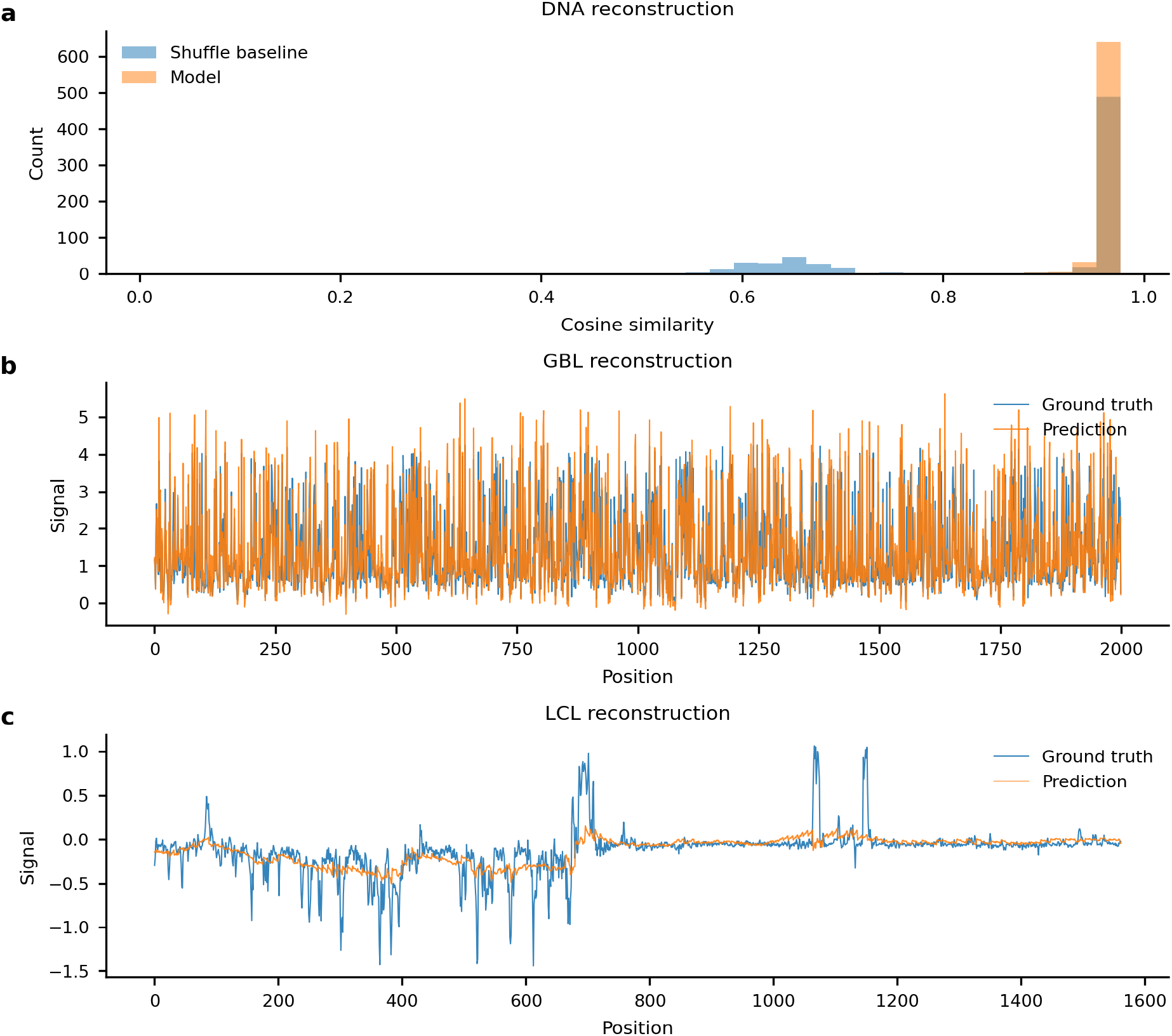
Modality-specific behavior of masked reconstruction. (a) DNA reconstruction was evaluated in embedding space using masked-token cosine similarity against a shuffled baseline, showing content-level recovery of masked DNA embedding tokens. (b) Representative global chromatin accessibility (GBL) reconstruction showing close agreement between predicted and ground-truth signal tracks. (c) Representative local chromatin accessibility (LCL) reconstruction after global normalization. LCL remained the most challenging modality: the model recovered part of the peak-related structure, but peak magnitude was still underestimated and local reconstruction errors remained.

We next examined how the masked reconstruction objective behaved across the three input modalities (Fig. 2). Because the modalities have different data structures, their reconstruction quality was assessed in modality-specific ways. DNA reconstruction was evaluated in embedding space, where masked-token cosine similarity exceeded a shuffled baseline, indicating content-level recovery of masked DNA embedding tokens. Additional token-level diagnostics further showed that reconstructed DNA tokens matched their corresponding masked positions and remained distinguishable from incorrect positions (Extended Data Fig. 2). Global chromatin accessibility showed stable track-level reconstruction, with predicted signals closely following the ground-truth profile. By contrast, local chromatin accessibility remained the most challenging modality: after global normalization, the model recovered part of the peak-related structure, but peak magnitude was still underestimated and local reconstruction errors remained.

These observations indicate that masked reconstruction learned useful information across modalities while also revealing the distinct difficulty of sparse local accessibility signals. DNA embeddings and GBL showed stable reconstruction, with mean cosine similarities of 0.963 and 0.972, respectively. LCL remained the most difficult modality, achieving a lower mean cosine similarity of 0.717 despite the use of global normalization. The baseline-improvement metric, defined as the ratio of model reconstruction score to a zero-prediction baseline averaged over three random seeds with masking ratios of 0.750 for DNA and GBL and 0.300 for LCL, further showed that DNA, GBL and LCL exceeded the zero-prediction baseline by 2.252-, 7.293-and 1.931-fold, respectively. These results indicate that the masked reconstruction objective effectively learned dense DNA-derived and global accessibility representations, while sparse local accessibility remained the main reconstruction bottleneck.

### Global normalization alleviates sparse-modality collapse

We next examined why LCL reconstruction was substantially more difficult than DNA embedding and GBL reconstruction. Local chromatin accessibility is sparse and peak-shaped: most positions are close to zero, whereas regulatory peaks occupy only a small fraction of each window. Under per-sample normalization, this distribution encourages a trivial near-zero solution during masked reconstruction, because predicting values close to zero can reduce reconstruction error for most masked positions without learning meaningful peak structure. This behavior explains the weak LCL improvement observed under the initial configuration and indicates that sparse local accessibility requires a normalization strategy that preserves peak contrast across windows.

To address this failure mode, we replaced per-sample normalization with global normalization for LCL. Instead of scaling each window independently, global normalization preserves the relative magnitude of accessibility signals across windows, reducing the incentive for the model to minimize reconstruction loss by predicting uniformly near-zero values. Diagnostic ablations under per-sample normalization tested whether LCL reconstruction could be improved by altering the LCL masking ratio, emphasizing peak-associated signals, adding an auxiliary peak-classification head or resampling high-accessibility windows. These variants produced only marginal LCL gains and, in several cases, reduced DNA or GBL reconstruction stability (Extended Data Table 1). In contrast, global LCL normalization substantially improved sparse-signal reconstruction, increasing the LCL improvement ratio to 1.931 and the LCL cosine similarity to 0.717. Although LCL reconstruction remained less accurate than DNA embedding and GBL reconstruction, global normalization shifted the model away from near-zero collapse and enabled partial recovery of local accessibility structure.

### Ablation analysis separates the effects of modality fusion and self-supervised training

We next performed ablation experiments to determine which components contributed to the downstream performance of DNA-MFM. Specifically, we compared DNA pooling, a DNA-only model, Random DNA-MFM and DNA-MFM to separate the effects of pooled DNA representations, encoder-based representation learning, multimodal fusion and self-supervised masked reconstruction. These configurations were evaluated across downstream regulatory-function tasks (Table 1), including activity classification, activity ranking, peak detection and peak-position prediction.

**Table 1.**
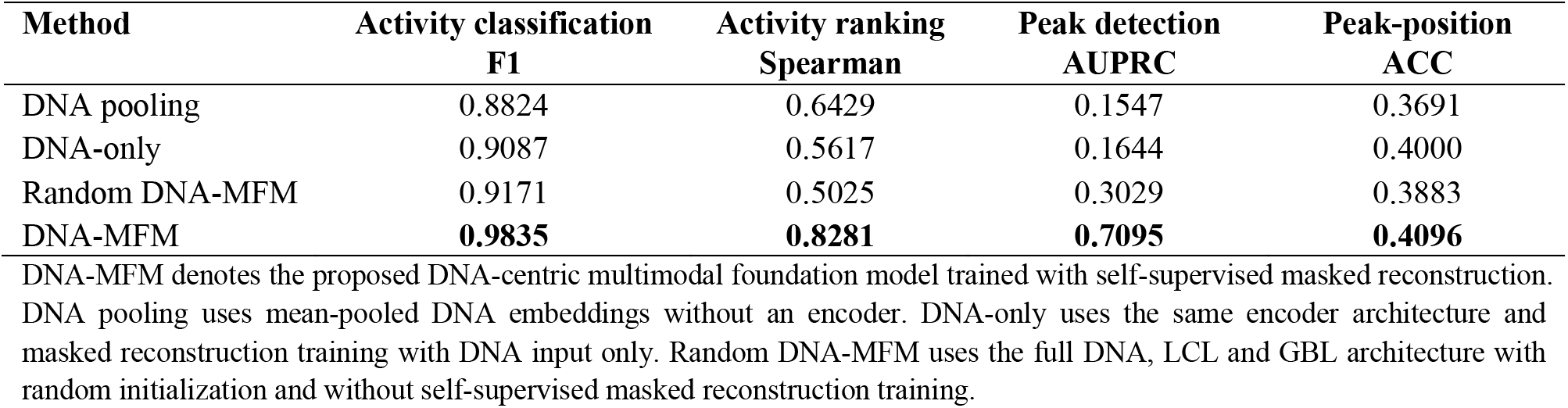
Modality and pre-training ablation across downstream tasks.

| Method | Activity classification | Activity ranking | Peak detection | Peak-position |
| --- | --- | --- | --- | --- |
|  | F1 | Spearman | AUPRC | ACC |
| DNA pooling | 0.8824 | 0.6429 | 0.1547 | 0.3691 |
| DNA-only | 0.9087 | 0.5617 | 0.1644 | 0.4000 |
| Random DNA-MFM | 0.9171 | 0.5025 | 0.3029 | 0.3883 |
| DNA-MFM | **0.9835** | **0.8281** | **0.7095** | **0.4096** |

DNA-MFM denotes the proposed DNA-centric multimodal foundation model trained with self-supervised masked reconstruction. DNA pooling uses mean-pooled DNA embeddings without an encoder. DNA-only uses the same encoder architecture and masked reconstruction training with DNA input only. Random DNA-MFM uses the full DNA, LCL and GBL architecture with random initialization and without self-supervised masked reconstruction training.

The ablation results show that DNA-MFM achieved the strongest performance across all four downstream tasks. DNA-only improved activity classification and peak-position prediction relative to DNA pooling, but did not improve activity ranking or peak detection, suggesting that encoder-based learning from DNA alone was insufficient for tasks requiring regulatory-state information. Random DNA-MFM improved peak detection compared with DNA-only, indicating that multimodal inputs and architectural capacity contributed to this task, but it remained substantially below DNA-MFM in activity ranking and peak detection. These results indicate that the downstream gains of DNA-MFM arise from the combination of multimodal fusion and self-supervised masked reconstruction, rather than from DNA features or model architecture alone. We further examined which design choices within the full DNA-MFM architecture were responsible for this performance gain (Table 2).

**Table 2.** Ablation of DNA-MFM design components across downstream tasks.

| Ablation setting | Activity classification | Activity ranking | Peak detection | Peak-position |
| --- | --- | --- | --- | --- |
|  | F1 | Spearman | AUPRC | ACC |
| DNA-MFM | **0.9835** | **0.8281** | **0.7095** | 0.4096 |
| VICReg | 0.9053 | 0.7412 | 0.2897 | 0.4298 |
| CLS pooling | 0.9149 | 0.7701 | 0.4895 | 0.4862 |
| All-token pooling | 0.8930 | 0.7121 | 0.2605 | 0.4521 |
| Per-sample LCL normalization | 0.9120 | 0.7491 | 0.3236 | 0.4404 |
| No masking curriculum | 0.9137 | 0.7274 | 0.1936 | 0.4478 |

All ablations use the full DNA-MFM architecture unless otherwise specified. The default DNA-MFM configuration uses global LCL normalization, mean-unmasked pooling, a masking curriculum and no VICReg regularization. Bold indicates the best value for the three primary metrics.

### DNA-MFM outperforms sequence-based embeddings across regulatory benchmarks

We next benchmarked DNA-MFM against sequence-based genomic embeddings to determine whether multimodal pre-training provided advantages beyond existing DNA-derived representations. We compared DNA-MFM with DNABERT-2, Nucleotide Transformer, Enformer tracks and DNA pooling under the same downstream evaluation protocol on held-out chromosomes (Table 3). This comparison assessed whether integrating chromatin accessibility improved regulatory-function prediction beyond sequence-only or sequence-derived embeddings.

**Table 3.** Benchmarking DNA-MFM against sequence-based genomic embeddings.

| Method | Input | Dimension | Activity classification | Activity ranking | Peak detection | Peak-position |
| --- | --- | --- | --- | --- | --- | --- |
|  |  |  | F1 | Spearman | AUPRC | ACC |
| DNABERT-2 | DNA | 768 | 0.6182 | 0.4248 | 0.1341 | 0.4351 |
| Nucleotide Transformer | DNA | 1024 | 0.7358 | 0.6299 | 0.1082 | 0.3979 |
| Enformer tracks | DNA | 5313 | 0.6606 | 0.8606 | 0.1898 | 0.4202 |
| DNA pooling | DNA | 1536 | 0.8824 | 0.6429 | 0.1547 | 0.3691 |
| DNA-MFM | DNA+ATAC | 256 | 0.9835 | 0.8281 | 0.7095 | 0.4096 |

All methods were evaluated on held-out chromosomes 20-22 using the same downstream evaluation protocol. Dimension denotes the embedding dimension used for downstream prediction. Enformer tracks denotes pooled Enformer-predicted functional tracks generated from DNA sequence. DNA+ATAC denotes DNA embeddings combined with local and global chromatin accessibility signals derived from ATAC-seq.

Benchmarking against sequence-based genomic embeddings showed that DNA-MFM provided the clearest gains in activity classification and peak detection. DNA-MFM achieved the highest F1 score and AUPRC among the compared embeddings, indicating improved performance on regulatory activity classification and chromatin peak detection. Activity ranking showed a different pattern, with Enformer tracks achieving the highest Spearman correlation, whereas DNA-MFM remained competitive. Peak-position prediction showed only small differences among methods. These results indicate that incorporating ATAC-derived chromatin accessibility most strongly improved regulatory activity classification and peak-associated signal detection. We further visualized the learned embedding space as a qualitative assessment of representation structure (Extended Data Fig. 3).

### DNA-MFM generalizes to external variant-effect datasets

We next evaluated whether DNA-MFM embeddings generalized beyond the internal regulatory-function benchmarks to external variant-effect datasets. We selected three datasets that represent distinct levels of variant-effect evidence: ClinVar non-coding variant classification for pathogenicity-related effects, GTEx eQTL effect-size prediction for expression-associated effects and PBMC caQTL effect-size prediction for chromatin-accessibility-associated effects. For these external analyses, Enformer tracks were used as a sequence-derived functional-track reference, and concatenated representations were evaluated to test whether DNA-MFM provided information complementary to these sequence-derived features. Together, these evaluations test whether the learned embeddings capture variant-associated functional signals across clinical, transcriptional and chromatin-accessibility contexts. We first evaluated DNA-MFM on ClinVar non-coding variant classification, comparing DNA-MFM with Enformer tracks and with a concatenated representation combining DNA-MFM and Enformer tracks (Table 4).

**Table 4.** External validation on ClinVar non-coding variants, GTEx eQTL and PBMC caQTL datasets.

| *ClinVar non-coding variant classification* |  |  |  |  |  |
| --- | --- | --- | --- | --- | --- |
| Representation | AUROC | AUPRC | F1 | ACC |  |
| Enformer tracks | 0.6141 | 0.3265 | 0.3656 | 0.6459 |  |
| DNA-MFM | 0.6829 | 0.3541 | 0.4140 | 0.6678 |  |
| DNA-MFM+Enformer tracks | 0.7012 | 0.4009 | 0.4360 | 0.6816 |  |
| *GTEx eQTL effect-size prediction* |  |  |  |  |  |
| Representation | Setting | n | Pearson | Spearman | R<sup>2</sup> |
| DNA-MFM | Region-level | 1,336 | 0.505 | 0.449 | 0.215 |
| DNA-MFM | Matched subset | 1,336 | 0.525 | 0.440 | 0.273 |
| Enformer $\Delta$ | All bins | 4,577 | 0.218 | 0.348 | -1.319 |
| DNA-MFM + Enformer $\Delta$ | All bins | 1,336 | 0.402 | 0.460 | -0.287 |
| *PBMC caQTL effect-size prediction* |  |  |  |  |  |
| Representation | Setting | n | Pearson | Spearman | R<sup>2</sup> |
| DNA-MFM | Region-level | 1,768 | 0.1401 | 0.1802 | -0.0416 |
| DNA-MFM | Matched subset | 1,768 | 0.3539 | 0.3764 | 0.1126 |
| Enformer $\Delta$ | All bins | 9,996 | 0.1748 | 0.5010 | -2.7090 |
| DNA-MFM + Enformer $\Delta$ | All bins | 1,768 | 0.2979 | 0.5771 | -1.2692 |

ClinVar: Enformer tracks denotes pooled Enformer-predicted functional tracks generated from DNA sequence. DNA-MFM + Enformer tracks denotes feature concatenation of the two representations. GTEx eQTL / PBMC caQTL: DNA-MFM denotes the multimodal embedding of the variant-containing window. Enformer $\Delta$ denotes the predicted change in Enformer outputs between alternative and reference alleles. DNA-MFM + Enformer $\Delta$ denotes feature concatenation before downstream regression. Region-level indicates assignment of the window-level embedding to each variant; matched subset denotes variants available for matched comparison. For Enformer-based rows, all-bin aggregation was used.

On ClinVar non-coding variant classification, DNA-MFM outperformed Enformer tracks across the main classification metrics, indicating that the multimodal embedding captured pathogenicity-related information beyond sequence-derived functional tracks. The concatenated representation of DNA-MFM and Enformer tracks achieved the strongest overall performance, suggesting that the two representations retained partially complementary information for non-coding variant classification. These results support the external utility of DNA-MFM embeddings for variant-effect prediction in a clinically relevant setting. We next evaluated expression-associated variant effects using GTEx eQTL effect-size prediction.

In GTEx eQTL effect-size prediction, DNA-MFM alone showed predictive signal for expression-associated variant effects, indicating that region-level multimodal embeddings captured information relevant to gene-regulatory variation. Enformer Δ provided a sequence-perturbation reference based on predicted changes between alternative and reference alleles. Compared with Enformer Δ, DNA-MFM showed stronger overall performance across the main regression metrics, whereas simple concatenation with Enformer Δ did not provide a consistent additional benefit. We finally evaluated chromatin-accessibility-associated variant effects using PBMC caQTL effect-size prediction, comparing DNA-MFM, Enformer Δ and their concatenated representation under the same regression framework.

In PBMC caQTL effect-size prediction, DNA-MFM showed predictive signal for chromatin-accessibility-associated variant effects. The matched DNA-MFM subset achieved the strongest Pearson correlation and R^2^, indicating that region-level multimodal embeddings captured information relevant to accessibility-linked regulatory variation. The concatenated representation achieved the highest Spearman correlation but did not improve Pearson correlation or R^2^, suggesting that simple feature concatenation provided metric-dependent rather than consistent gains. Together with the GTEx eQTL results, these findings indicate that DNA-MFM embeddings generalize to external variant-effect datasets involving both gene expression and chromatin accessibility.

Together, the three external validations show that DNA-MFM provides useful standalone representations for variant-effect analysis across pathogenicity-related classification, expression-associated eQTL prediction and chromatin-accessibility-associated caQTL prediction. DNA-MFM improved ClinVar classification relative to Enformer tracks and showed stronger Pearson and R^2^ performance than Enformer Δ in both eQTL and caQTL effect-size prediction. Feature concatenation with Enformer-derived features provided metric-dependent benefits, but did not consistently improve all regression metrics. These findings support DNA-MFM as a reusable region-level multimodal representation for external functional and variant-effect analyses.

## Discussion

We developed DNA-MFM, a self-supervised DNA-centric multimodal foundation model that integrates DNA sequence embeddings with local and global chromatin accessibility signals to learn reusable genomic-region representations. The results show that ATAC-derived accessibility information strengthens representation learning for regulatory tasks, especially activity classification and chromatin peak detection, and provides useful external signal for ClinVar, GTEx eQTL and PBMC caQTL analyses. These findings support the value of DNA-centered representations that incorporate regulatory-state information alongside sequence-derived features.

A major challenge in this setting is the mismatch between dense DNA embeddings and sparse, peak-shaped accessibility signals. Under per-sample normalization, local chromatin accessibility showed weak reconstruction consistent with sparse-modality collapse. Global normalization substantially improved LCL reconstruction and supported stable multimodal training, although LCL remained the most difficult modality. This result highlights normalization as a key component for aligning heterogeneous genomic modalities during self-supervised pre-training.

The downstream and external evaluations further define the role of DNA-MFM. DNA-MFM provides a reusable region-level multimodal embedding that captures regulatory context, while Enformer-derived perturbation features provide allele-specific sequence-effect information. In eQTL and caQTL analyses, feature concatenation produced metric-dependent gains, indicating that region-level regulatory context and allele-specific perturbation signals may benefit from more structured integration in future models. The current framework focuses on DNA sequence and ATAC-derived chromatin accessibility, and can be extended to additional regulatory modalities such as histone modifications, DNA methylation and expression-associated signals.

## Methods

### Dataset and preprocessing

We constructed the DNA-MFM training and evaluation datasets from genome-wide windows paired with DNA sequence-derived embeddings and ATAC-seq^21^ chromatin accessibility signals. Data were derived from the EnformerCelltyping framework^10^, using the human reference genome (hg38) and matched chromatin accessibility tracks. Genomic windows were sampled genome-wide with a sliding-window strategy, and each window spanned approximately 114 kb aligned to the Enformer output resolution. Window-level features were pre-computed before model training to ensure consistent input representation across pre-training and downstream evaluation.

Chromosomes 1-19 were used for pre-training, yielding approximately 20,000 genomic windows. Chromosomes 20-22 were held out for evaluation, yielding approximately 940 windows. This chromosome-level split was used to prevent sequence-level leakage between training and evaluation.

Each genomic window was represented by three input streams. DNA embeddings were extracted from raw DNA sequence using a pre-trained chopped Enformer model, producing 1,536-dimensional continuous embeddings across 896 positions. Local chromatin accessibility (LCL) was represented as a one-dimensional ATAC-seq signal track at 128-bp resolution. Global chromatin accessibility (GBL) was represented as a broader one-dimensional accessibility track over the same genomic context.

To reduce the influence of near-zero background windows, we applied coverage-aware filtering during window construction. Candidate windows were evaluated using chromatin-accessibility coverage, based on the fraction of non-zero bins and peak-containing signal within the LCL or GBL tracks. Windows with insufficient accessibility coverage were removed, and approximately 72% of candidate windows were retained after filtering. This step reduced the dominance of uninformative background windows while preserving the chromosome-level train–evaluation split (Extended Data Fig. 1).

### Model architecture

To convert heterogeneous genomic inputs into a common token representation, we used modality-specific normalization and projection for DNA, LCL and GBL. DNA embeddings were treated as position-wise tokens and projected from 1,536 to 256 dimensions while preserving the 896-position sequence length. LCL values were normalized using global statistics estimated from the training windows, preserving cross-window differences in accessibility magnitude and reducing near-zero collapse of sparse local signals. The normalized LCL track was divided into non-overlapping patches with a patch size of 8, and each patch was projected to a 256-dimensional token. GBL signals were robustly normalized per sample to reduce the influence of extreme values within each global accessibility track, and were divided into non-overlapping patches with a patch size of 16, yielding 912 patch tokens after projection to 256 dimensions.

The DNA, LCL and GBL token streams were concatenated with a learnable CLS token and combined with learnable positional embeddings and modality-type embeddings. Positional embeddings preserved token order, whereas modality-type embeddings distinguished the CLS, DNA, LCL and GBL streams before input to the shared Transformer encoder.

The concatenated multimodal token sequence was encoded using a shared Transformer encoder with bidirectional self-attention^22^, allowing DNA, LCL and GBL tokens from the same genomic window to exchange information in a common latent space. The encoder used six layers, a hidden dimension of 256, eight attention heads, a feed-forward dimension of 1,024, GELU activation, dropout of 0.1 and a pre-norm layout. The final window-level representation was obtained by mean pooling over unmasked tokens from the final encoder layer, producing a 256-dimensional embedding for each genomic window.

### Self-supervised training

The model was trained with a self-supervised masked reconstruction objective^11^ over the three modality-specific token streams. During training, masking positions were sampled independently within each modality, with masking ratios of 0.75 for DNA, 0.30 for LCL and 0.75 for GBL. At masked positions, the original token values were replaced with a learned placeholder vector, and the model was optimized to reconstruct the original token values using the unmasked DNA, LCL and GBL tokens from the same genomic window. The lower masking ratio for LCL was used to account for the sparsity of local accessibility signals. Reconstruction was optimized using modality-weighted Huber losses computed only at masked positions, balancing reconstruction across high-dimensional DNA embedding tokens and lower-dimensional accessibility patch tokens.

Masked reconstruction quality was evaluated by comparing predicted and ground-truth values at masked positions for DNA embeddings, LCL and GBL. For each modality, reconstruction performance was summarized using mean cosine similarity, median cosine similarity and baseline improvement, defined as the ratio between the model reconstruction score and the corresponding zero-prediction baseline score. Additional token-level reconstruction analyses were performed for DNA embeddings (Extended Data Fig. 2).

Training used AdamW^13^ optimization. The learning rate was linearly warmed up during the initial training phase and then decayed with a cosine schedule. A masking curriculum was applied during the warmup period, increasing the masking ratio from an initial value to the modality-specific target ratios. After training, the encoder was used to produce fixed 256-dimensional window-level embeddings for subsequent analyses. Training hyperparameters are summarized separately (Supplementary Table 1).

### Downstream regulatory-function evaluation

Downstream regulatory-function evaluation was performed on chromosomes 20-22 using linear probes on fixed window-level embeddings. Task labels were derived from six Enformer-resolution histone-mark tracks from the Enformer Celltyping framework: H3K27ac, H3K4me1, H3K4me3, H3K9me3, H3K27me3 and H3K36me3. These labels were used to evaluate regulatory activity from complementary perspectives, including signal presence, signal strength, peak detection and peak localization.

For regulatory activity classification, binary labels were derived from the top and bottom quantiles of the log1p-transformed mean histone-mark signal, using q = 0.1 and q = 0.9. Classification was performed with L2-regularized logistic regression. For regulatory activity ranking, continuous histone-mark signal values were used as regression targets and evaluated with ridge regression. For chromatin accessibility peak detection, peak-calling labels derived from LCL signals were used as positive labels. For peak-position prediction, the relative peak position within each window was classified into left, centre or right. The primary metrics were F1 score for activity classification, Spearman correlation for activity ranking, AUPRC for peak detection and top-1 accuracy for peak-position prediction. All probes used a regularization strength of 1.0.

### External variant-effect evaluation

External variant-effect evaluation was designed to test whether DNA-MFM embeddings generalized from internal regulatory-function benchmarks to independent variant-effect settings. Three datasets were used to cover complementary functional contexts: ClinVar non-coding variants for pathogenicity-related effects, GTEx eQTL variants for expression-associated effects and PBMC caQTL variants for chromatin-accessibility-associated effects.

For ClinVar evaluation, pathogenicity labels from the 2024 ClinVar release^15^ were converted into a binary classification task by merging Pathogenic and Likely pathogenic variants into the pathogenic class and Benign and Likely benign variants into the benign class. Variants with conflicting interpretations or uncertain significance were removed. To focus on regulatory variant effects, only non-coding variants in 5′UTR, 3′UTR, intronic and intergenic regions were retained. Each variant was assigned the DNA-MFM embedding of the genomic window containing the variant, and logistic regression with L2 regularization was used for pathogenicity classification. ClinVar performance was evaluated using AUROC, AUPRC, F1 score and accuracy.

For GTEx eQTL evaluation, significant eQTLs from GTEx v8 Whole Blood^14^ were used to assess expression-associated variant effects, with normalized effect size used as the regression target. Each variant was mapped to the genomic window containing it, and the corresponding DNA-MFM embedding was used as a region-level representation. Ridge regression with L2 regularization was used for effect-size prediction. As a sequence-based reference, Enformer Δ features were generated for SNPs by predicting functional tracks from reference and alternate allele sequences and taking the difference between alternate and reference predictions. DNA-MFM features, Enformer Δ features and their concatenation on the overlap subset were evaluated using Pearson correlation, Spearman correlation and R^2^.

For PBMC caQTL evaluation, chromatin-accessibility-associated effect sizes were assessed using a public PBMC single-cell ATAC-seq caQTL dataset^16^. Each caQTL variant was mapped to its corresponding genomic window, and the DNA-MFM embedding of that window was used as a region-level representation for ridge regression. Enformer Δ features and concatenated DNA-MFM plus Enformer Δ features were evaluated as comparison settings. Performance was measured using Pearson correlation, Spearman correlation and R^2^.

### Ablation settings

To separate the effects of DNA-derived representations, multimodal fusion and self-supervised training, we evaluated four representation settings under the same downstream regulatory-function protocol. DNA pooling used mean-pooled DNA embeddings without an additional encoder. DNA-only used the same Transformer encoder architecture with DNA input only. Random DNA-MFM used the full DNA, LCL and GBL input architecture with random initialization and without self-supervised masked reconstruction training. DNA-MFM used the full multimodal architecture after self-supervised masked reconstruction training.

Additional ablation settings were evaluated within the DNA-MFM framework to examine the contribution of key design choices, including pooling strategy, LCL normalization, masking curriculum and VICReg^12^ regularization. Sparse-LCL reconstruction variants, including adjusted LCL masking ratios, increased LCL reconstruction weight and peak-aware reconstruction objectives, were compared with the final global-normalization configuration. Hyperparameter sensitivity was further assessed across model and training settings (Extended Data Fig. 4).

## Data availability

Genomic sequence and chromatin accessibility data used in this study are publicly available through the EnformerCelltyping project (https://github.com/neurogenomics/EnformerCelltyping) and the UCSC Genome Browser hg38 reference genome (https://genome.ucsc.edu). ClinVar data from the 2024 release were obtained from the NCBI ClinVar database (https://www.ncbi.nlm.nih.gov/clinvar/). GTEx v8 eQTL data were obtained from the GTEx Portal (https://gtexportal.org). Processed genomic-window metadata and derived DNA-MFM embeddings generated in this study will be made available upon publication.

## Code availability

Custom code for model training, embedding generation and downstream evaluation is available at https://github.com/Yukyin/dna-mfm. The repository includes all scripts required to reproduce the main results reported in this manuscript.

## Acknowledgements

The author thanks the developers of the public genomic resources and software tools used in this study.

## Author contributions

Y.C. conceived the study, designed the model, performed the experiments, analyzed the data and wrote the manuscript.

## Extended Data Figures

**Extended Data Fig. 1.**
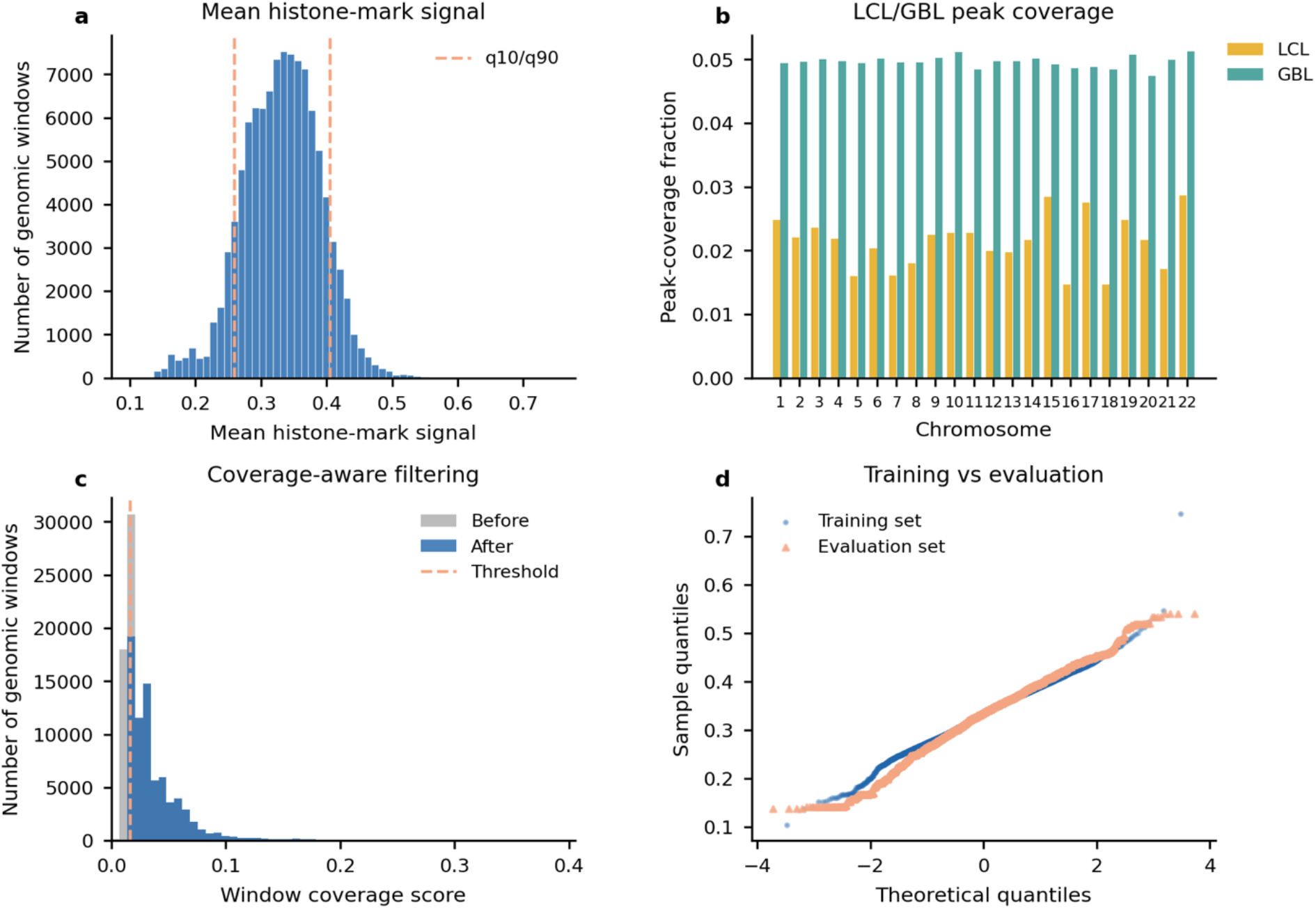
Training-window distribution and quality control. (a) Distribution of mean histone-mark signal across genomic windows. (b) LCL and GBL peak-coverage distributions across chromosomes. (c) Number of genomic windows before and after coverage-aware filtering. (d) Quantile–quantile plots comparing feature distributions between training and evaluation chromosome sets.

**Extended Data Fig. 2.**
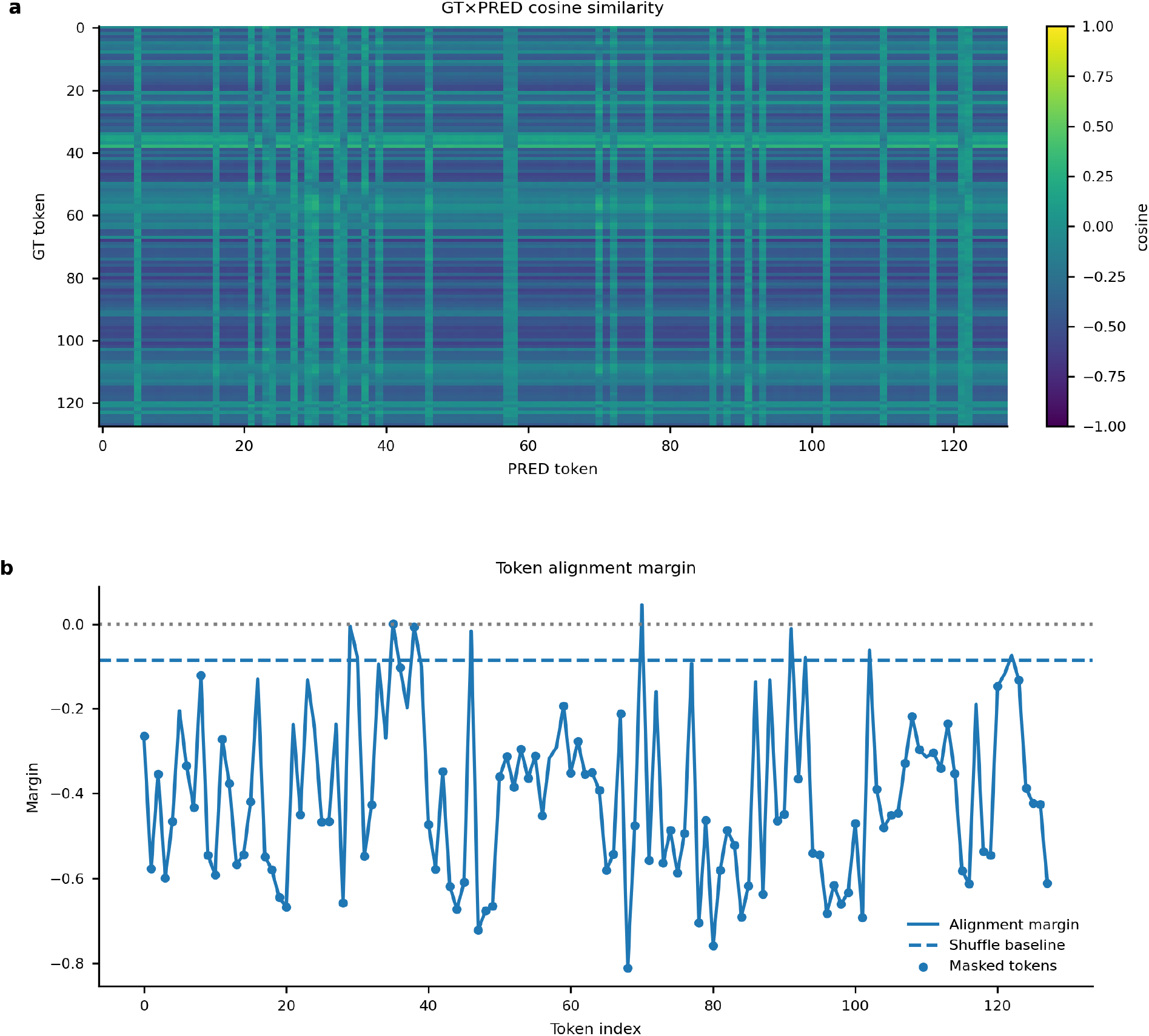
Detailed analysis of DNA token reconstruction. (a) Token-by-token cosine similarity matrix for a representative test window. (b) Alignment margin between the correct masked position and the best-matching incorrect position.

**Extended Data Fig. 3.**
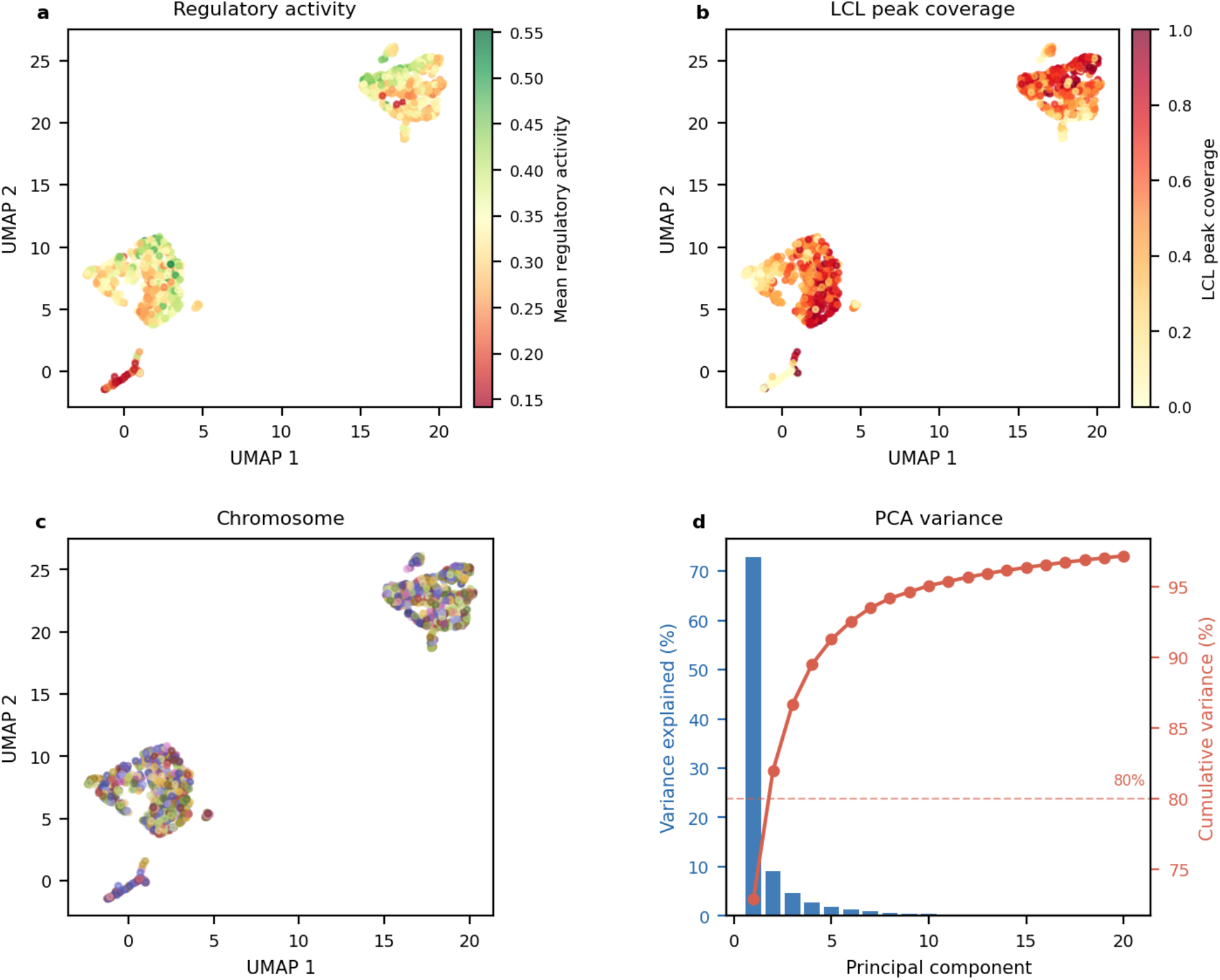
Embedding-space visualization. (a) UMAP projection of 1,000 randomly sampled training-window embeddings coloured by mean regulatory activity. (b) The same UMAP projection coloured by LCL peak coverage. (c) The same UMAP projection coloured by source chromosome. (d) Variance explained by the top 20 principal components of the full 20,000-window embedding library.

**Extended Data Fig. 4.**
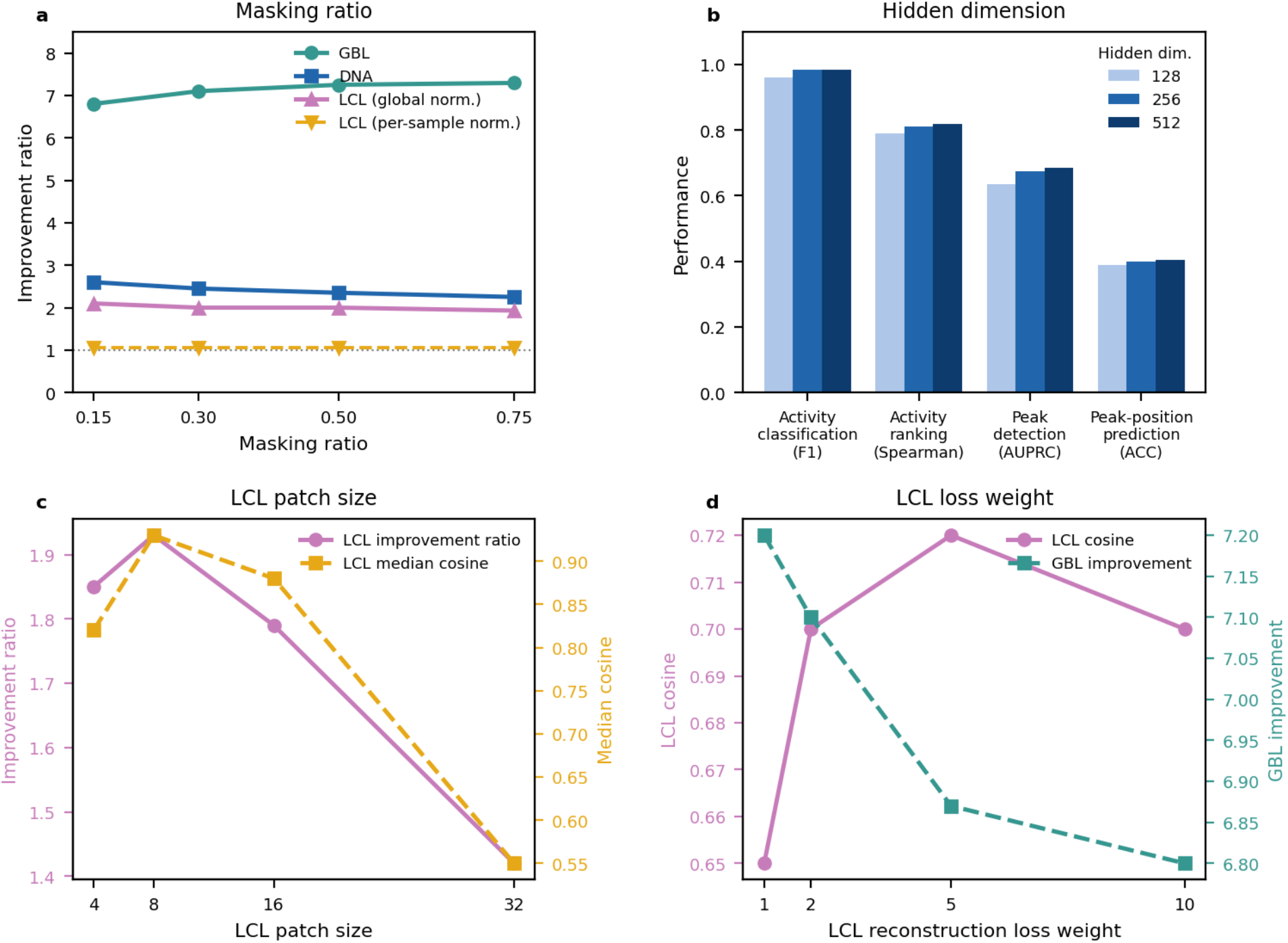
Hyperparameter sensitivity of DNA-MFM. (a) Effect of masking ratio on reconstruction improvement ratio for each modality. (b) Effect of encoder hidden dimension on downstream task performance. (c) Effect of LCL patch size on LCL reconstruction improvement ratio. (d) Sensitivity of LCL reconstruction performance to the LCL reconstruction loss weight.

## Extended Data Tables

**Extended Data Table 1.** Diagnostic ablations for sparse LCL reconstruction.

| Ablation setting | Normalization | LCL<br>improvement<br>ratio | LCL<br>cosine<br>similarity | DNA<br>improvement<br>ratio | GBL<br>improvement<br>ratio |
| --- | --- | --- | --- | --- | --- |
| LCL masking ratio 0.15 | Per-sample | 1.046 | 0.247 | 2.128 | 6.171 |
| LCL masking ratio 0.20 | Per-sample | 1.041 | 0.246 | 2.132 | 5.952 |
| LCL masking ratio 0.30 | Per-sample | 1.104 | 0.334 | 2.131 | 5.898 |
| LCL masking ratio 0.40 | Per-sample | 1.074 | 0.270 | 2.135 | 5.907 |
| Peak-aware weighted loss | Per-sample | 1.090 | 0.332 | 2.129 | 3.566 |
| Auxiliary peak-classification head | Per-sample | 1.043 | 0.247 | 2.011 | 2.989 |
| Top-30% accessibility resampling | Per-sample | 1.039 | 0.237 | 1.965 | 3.326 |
| Global LCL normalization | Global | 1.931 | 0.717 | 2.252 | 7.931 |

Improvement ratios were calculated relative to the zero-prediction baseline. All settings used an LCL reconstruction weight of 5 and an LCL patch size of 8.

## Supplementary Tables

**Supplementary Table 1.** Training hyperparameters for DNA-MFM.

| Hyperparameter | Value |
| --- | --- |
| Optimizer | AdamW ($\beta_1 = 0.9$, $\beta_2 = 0.999$) |
| Learning rate | 2e-4 |
| Weight decay | 0.05 |
| Batch size | 32 |
| Training epochs | 10 |
| Training iterations | 33,060 |
| Warmup iterations | 9,918 |
| Initial masking ratio | 0.15 |
| Random seeds | 42, 43, 44 |
| GPU | A100 40G |

